# The native - exotic plant choice in green roof design: using a multicriteria decision framework to foster urban biodiversity

**DOI:** 10.1101/2022.01.07.475351

**Authors:** Ana A. Calviño, Julia Tavella, Hernán M. Beccacece, Elizabet L. Estallo, Diego Fabián, María Laura Moreno, Adriana Salvo, María Silvina Fenoglio

## Abstract

Green roofs are considered key elements of the urban green infrastructure since they offer several environmental benefits, including habitat provision for arthropods. To achieve these benefits and ensure green roof success, an appropriate plant selection is an important step in the design of these infrastructures, especially where green roof technology is emerging like in South American cities. So far, decisions of using native or exotic plant species in green roofs had never been evaluated taking into account the plant potential to foster beneficial arthropods. By applying an integrative multicriteria decision framework that combined the habitat template hypothesis with the potential of plants to attract floral visitors and natural enemies, we obtained a ranked set of candidate native and exotic plant species. Among the best-ranked candidate species, we further compared the performance of six native and six exotic species in 30 experimental green roofs installed in Córdoba city, Argentina. To evaluate plant success, the occurrence and cover of each species were recorded one year after establishment under two management conditions: regular watering and weeding of spontaneous plants, and no management (15 roofs each). All selected species increased their vegetative cover one year after establishment. More interestingly, native plants had an advantage over exotic plant species as they exhibited a significantly higher occurrence and a slightly higher cover with no management than exotics. Native annuals were able to reseed the following season even in the absence of management, thus highlighting the relative importance of lifespan as a useful plant trait for future studies in green roof design. Given that green roofs are one of the possible solutions to ameliorate the negative effects of urban habitat loss on arthropod diversity, the development of an integrative multicriteria decision framework that takes into account the potential of native and exotic plant species for promoting beneficial arthropods would give a new twist in plant selection processes for green roofs.

## 1. Introduction

Green roofs are considered key elements of the urban green infrastructure as they contribute to runoff control, carbon sequestration, temperature regulation and habitat or food provision for different organisms, mostly arthropods (MacIvor and Ksiazek, 2015; Thuring and Grant, 2016; Guarino et al., 2021; Heim et al., 2021). To achieve all these environmental benefits and ensure green roof success, an appropriate plant selection is an important step in the design of these infrastructures. Until now, decisions regarding plant species’ origin in green roofs have been evaluated in relation to its roof adaptability but not including the plants’ potential to foster beneficial arthropods in an integrative multicriteria decision framework. Moreover in South American cities, where green roof technology and especially the selection and use of native plant vegetation are still in its infancy (but see Jaramillo Pazmino, 2016; Cáceres et al., 2018), the use of decision tools would be helpful to integrate previous knowledge on this matter with novel conservation goals.

At the beginning of their history, green roofs were thought of as fire protection covers so then spontaneous plant species started to colonize them (Dabija 2019). Nowadays, decisions regarding which species are most suitable for green roofs encompass a diverse universe of criteria. Given that rooftops are particularly harsh environments, the selection of plant species was initially based on the use of plant traits as hardiness surrogates. Accordingly, drought-tolerant succulent plant species, well adapted to the stressful conditions of the roof, were primarily chosen. Among these, and mostly out of its native range, *Sedum* (Crassulaceae) species usually dominate green roof vegetation over the world (Cook-Patton, 2015). Several advantages have been found in *Sedum* species, ranging from their high survival rate to their temperature regulation and water retention capabilities (Butler and Orians, 2011), in contrast with their limited value for urban biodiversity (Kiehl et al., 2021). Well beyond the widely used *Sedum* species, nevertheless, the trait-based selection framework significantly contributed to improving the quality of the decisions around plant selection, broadening the ecosystem services provided by green roofs (Lundholm and Walker, 2018; Heim et al., 2021). Most certainly, a qualitative leap in the history of modern green roof design came with the introduction of the habitat template hypothesis into the plant selection process (Lundholm, 2006). By taking into account a habitat analog, the habitat template hypothesis states that natural habitats with similar abiotic characteristics to roofs provide reliable information about the potential plant species to be used. In fact, there are several successful experiments that, by assuming habitat templates, have arrived at a plant species pool able to succeed in green roofs (e.g., Kiehl et al., 2021; Ksiazek-Mikenas et al., 2021). Thus, this approach provides an optimum ecological framework for selecting plant species which, in addition, may be easily integrated with trait-based approaches (e.g., Lundholm and Walker, 2018).

Regarding the relative success of exotic versus native plant roof cover, most examples are from the northern hemisphere with no clear performance advantages of any group (Butler et al., 2012). For its part, other temperate, semi-arid and arid regions of the world may provide good candidates of native species other than the traditional *Sedum* vegetation roofs’ cover (e.g., Cáceres et al., 2018; Yee et al., 2021), but the promising horizon of better native alternatives remains to be tested within a common comparative framework. This is crucial to address the relative value of a given plant on the basis of its origin. In cases where the exotic vs. native species pools are selected by different criteria (i.e., exotics chosen by their use in roofs but natives by a habitat analog), the origin effect may lose strength as well as the conclusions obtained could gain inconsistency.

According to the “adaptation argument”, native plant species would perform better than exotics in their native habitat, as they use water more efficiently than their non-native counterparts (Butler et al., 2012). Native plant species are, in addition, suitable for extensive green roofs (Cascone, 2019) which are characterized by low maintenance vegetation able to self-sow, traits usually satisfied by local native flora (Sutton, 2015). Moreover, evidence from green roofs sustains a greater potential of native over exotic plant species to promote and support native biodiversity (Cook-Patton, 2015; Khiel et al., 2021). Given that green roofs can support a considerable diversity of arthropods from several functional groups (e.g., Knapp et al., 2019; Fabián et al., 2021), it is expected that the use of local native plant species will favour the urban native arthropod fauna such as herbivores, pollinators and parasitoids (e.g., Mata et al., 2021). In spite of this, the potential of plants to attract beneficial arthropods, whether being native or not, has never been taken into account when selecting plants for roofs.

All these aspects highlight the need to integrate the traditional decision frameworks designed to select plant species able to survive in extensive green roofs, like habitat template analogs, to ecological plant attributes relevant for the co-occurring urban fauna. However, neither the habitat template approach has been considered to foster biodiversity at higher trophic levels (Ksiazek-Mikenas et al., 2021), nor the comparison regarding plants’ performance has been yet addressed after applying the same selection framework to both native and exotic plant species.

Multicriteria decision-making analysis (MCDA) is a useful approach to deal with complex human decisions and a strong tool to “validate our thinking” by weighting our previous knowledge about a given problem (Saaty, 2004). In addition, it has provided good examples of how to resolve complex decisions regarding green infrastructure planning and design (e.g., Asgarzadeh et al., 2014; Vlachokostas et al., 2014; Rosasco and Perini, 2019) and conservation issues (Adem Esmail and Geneletti, 2018). Here, we employed MCDA to rank and then select six native and six exotic plant species which were established in 30 experimental green roofs in Córdoba city, Argentina, as a part of a larger project designed to test the effect of plant origin on arthropod diversity. By combining the habitat template hypothesis with surrogates of plant affinity for beneficial arthropods in a multicriteria decision framework, we obtained a ranked set of candidate native and exotic plant species expected to tolerate roof conditions and able to attract floral visitors and natural enemies. In turn, and in order to have a measure of plant success, the occurrence and cover of each species were recorded one year after establishment under two management conditions: green roofs i) with regular watering and weeding of spontaneous vegetation, and ii) without management (i.e., extensive green roof). Based on the adaptation argument (Butler et al., 2012) we expect that native plant species will perform better than exotics given that the former requires less maintenance and water.

## 2. Methods

### 2.1 Species lists

The species selection procedure is summarized in Figure 1. First, an initial list of potential plant species for green roofs was built on the basis of published lists of plant species already registered in green roofs all over the world. To do so, we performed an initial literature search, with the keywords “green”, “roof” and “plant” in the Google Scholar platform. After that we selected a pool of 29 published articles based on the following criteria: 1) they should provide information on plant species registered as growing in green roofs either as spontaneously or cultivated; 2) papers with only lists of recommended plant species but not tested in green roofs were discarded, 3) priority was given to studies that provide any measure of the plants’ performance in the roofs (i.e., relative frequency, cover, density, etc.). After applying those criteria, we obtained an initial plant list with a total of 1393 plant species, representing green roofs from Europe, Asia, North, and South America. Second, the list was refined on the basis of plant life form (only herbaceous plants were included), and then regarding their occurrence in the Argentinian flora website (www.floraargentina.edu.ar) either as native or not, or their citation in any of the ornamental and cultivated plants’ guides from Dimitri and Parodi (1977) and Hurrell et al. (2006, 2007, 2009, 2017). In addition, and for ornamental exotic species only, we checked their availability in wholesale local nurseries to ensure that those species will be able to be reproduced in the short term. The final plant list contained 117 candidate species that were classified as native with a political criterion of nativeness (*sensu* Berthon et al., 2021). Accordingly, a species was considered as native whether it was classified as such in the Argentinian flora and registered in Córdoba province. As a result, we obtained 57 native and 60 exotic candidate species in the final plant list (Figure 1, Supplementary Material Table S1).

**Figure 1.**
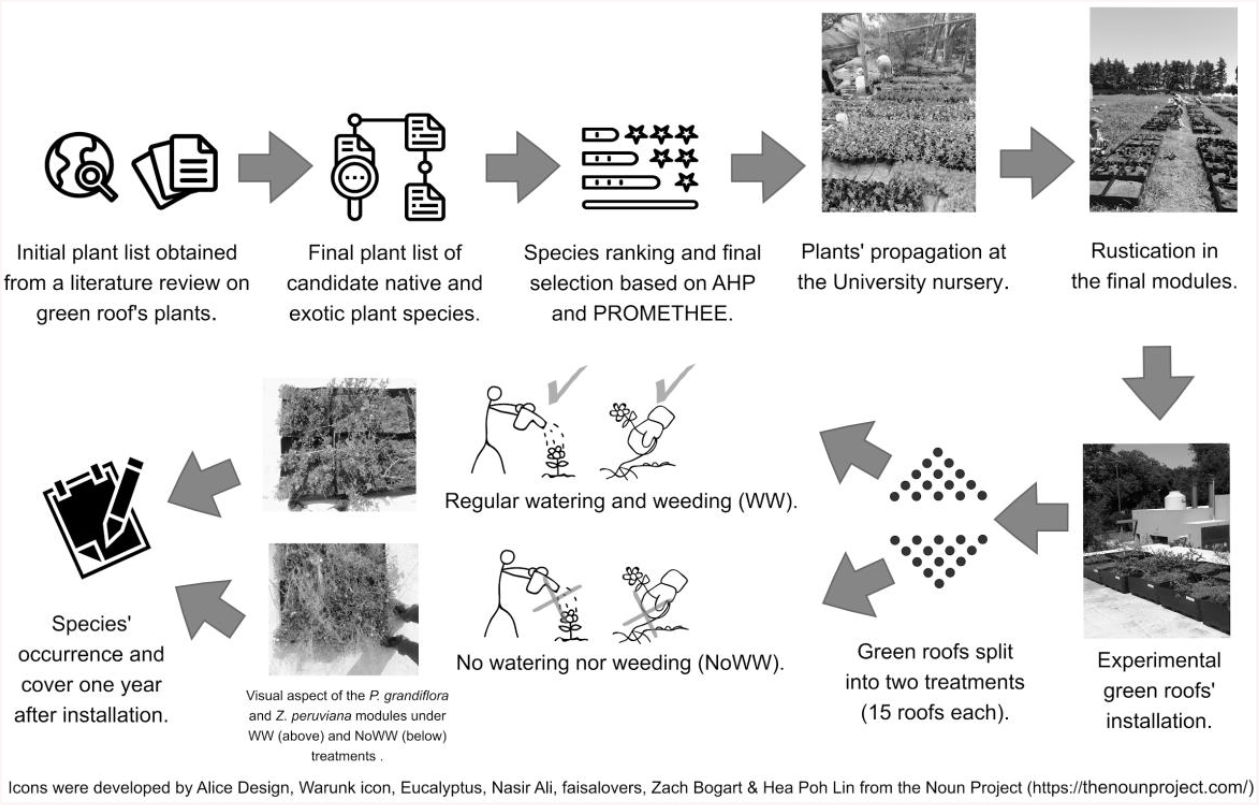
Flow diagram showing the steps followed for the selection of native and exotic plant species to be installed in the experimental green roofs in Cordoba city, Argentina. AHP: Analytical Hierarchical Process, PROMETHEE: Preference Ranking Organization Method of Enrichment Evaluation. Please, see Methods for further details.

### 2.2 Multicriteria decision making analyses

#### 2.2.1 The decision model

Under the generic designation of Multicriteria Decision Making Analysis or aiding process (MCDA) there is a diverse group of systematic approaches originally designed to deal with multiple and often conflicting alternatives within a common decision framework (Marttunen et al., 2017). Here, to rank the 117 candidate plant species (57 natives and 60 exotics), we combined two decision-making tools that use pairwise comparisons between alternatives (i.e., plant species). One procedure, the Analytical Hierarchical Process (AHP; Saaty, 1980) was only used here to define and weight the criteria that plant species should ideally meet to succeed in green roofs and have the potential for attracting beneficial arthropods.

A second procedure, the Preference Ranking Organization Method of Enrichment Evaluation (PROMETHEE; Brans et al., 1986) was used to rank the species according to the weight of the criteria established previously by the AHP. The combination of these two procedures is sustained by the fact that AHP gives an accurate estimate to weight the selection criteria, whereas PROMETHEE is preferred over other MCDA tools for decision problems involving few criteria and many decision alternatives (Si et al., 2016). An AHP usually starts with a graphical representation of the goal and the principal and subordinate criteria used in the decision (Figure 2). We used two types of decision criteria: one group of criteria to define the potential of a given plant species to tolerate green roof conditions, and the other group of criteria to infer the potential of a given plant species to attract flower visitors and natural enemies (Figure 2).

**Figure 2.**
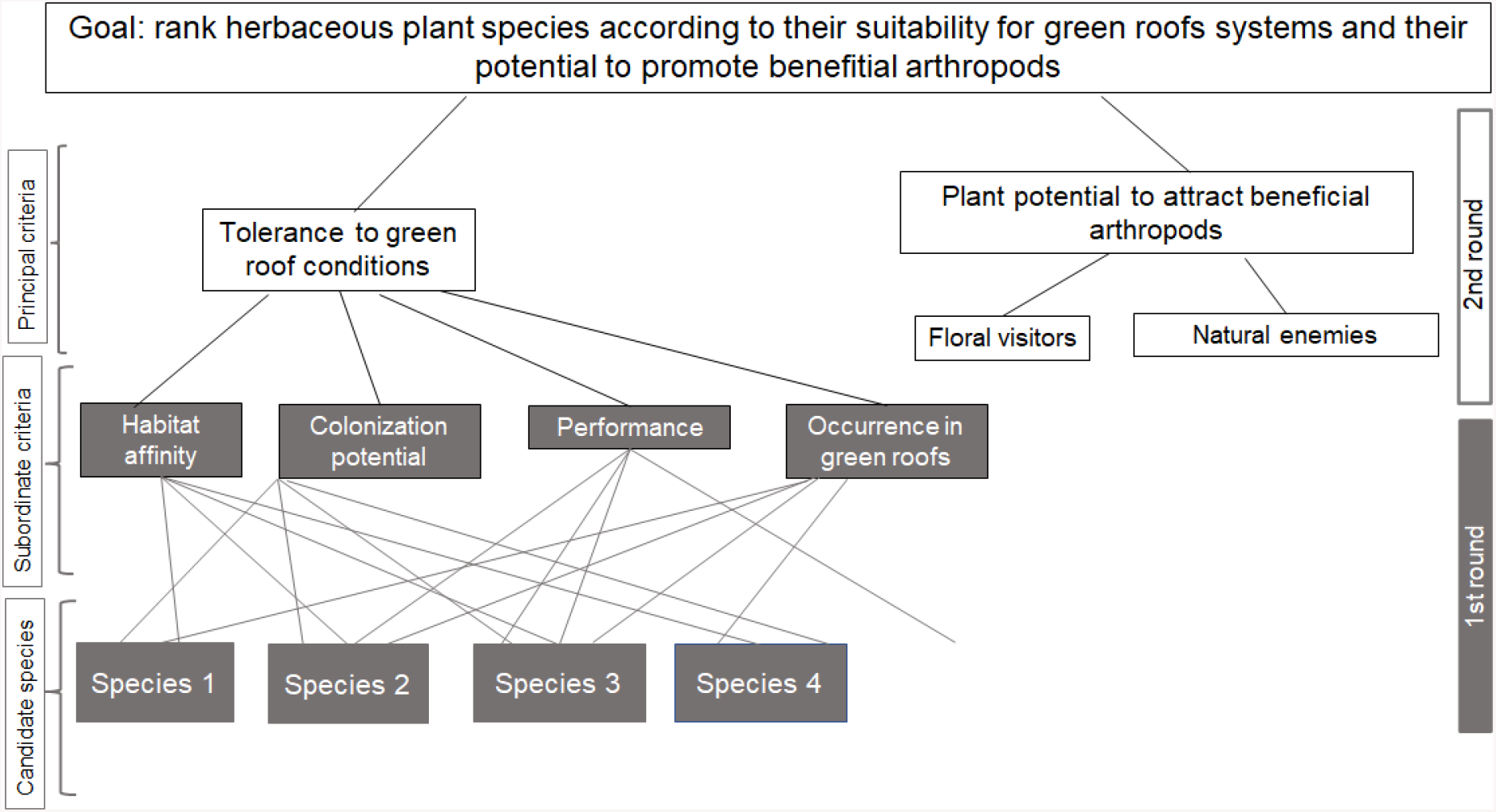
Structure of the criteria employed in the Analytical hierarchical process (AHP). Subordinate and principal criteria were used in the first (grey) and second (white) decision rounds, respectively. Candidate species were the native and exotic species included in the final plant list Supplementary Material Table S1). Rankings of species were obtained with the Preference Ranking Organization Method of Enrichment Evaluation (PROMETHEE) on the basis of the criteria weights obtained by AHP. Please, see methods for further details.

#### 2.2.2 Outranking procedure and criteria definition

The complete outranking process was performed in two decision rounds, every round consisting of an AHP and a PROMETHEE procedure for native and exotic candidate species, separately. The output of the first round gave us the species’ ranking only according to their tolerance to green roof conditions and based on four criteria: habitat affinity, colonization potential, performance, and occurrence (Figure 2). To define each criterion’s value, we performed additional literature searches to fulfil the necessary information for each plant species. References used to support the criteria value for each candidate species are provided at the Supplementary Material Reference List S1-S2. Habitat affinity was defined as the theoretical similarity of rooftops with natural habitats: rock outcrop habitats (including stonecrops, cliffs), ruderal (i.e., roadside, escaped plants), and other habitats (e.g., sand dunes, grasslands). A given plant species may spontaneously occur in more than one habitat, and so we gave maximal value to species recognized as living in both rocky and ruderal habitats. By doing so, we aimed to prioritize the habitat plasticity of a given species and its potential ability to cope with a wider range of environmental features. For instance, rocky habitats may share some features with green roofs but differ in others such as typically shallow soil depths of green roofs (Lundholm, 2006). In addition, ruderal habitats are considered good surrogates for extensive green roofs (Cascone, 2019) and they have been shown to favour the functional diversity of insects (e.g., Kalarus et al., 2019) so that species occurring both in rocky and ruderal habitats may combine highly preferred traits according to our goal (Figure 2). Colonization potential was a binary parameter used to identify those plant species already registered as spontaneous in green roofs or not in literature. Performance was also a categorical criterion gathering the information regarding the species’ cover, germination, or survival registered in the green roofs. Finally, occurrence represented the number of studies that cite the presence of a species, always considering different green roof studies

The second decision round gave us the final species’ ranking on the basis of three criteria: the plant potential to tolerate green roof conditions (obtained by the first decision round and equivalent to the previous species’ rank order number), and two criteria defined as relevant for promoting beneficial arthropods: the potential of plants to attract both floral visitors and natural enemies (Figure 2). Attractiveness to each target group was defined as the number of arthropod taxa (i.e., orders) registered for each plant species. For floral visitors (Hymenoptera, Diptera, Coleoptera, and Lepidoptera) we considered the total number of orders registered in the literature for each plant species. For natural enemies, we counted the number of recorded phytophagous orders cited in the literature for each plant species, considering then the number of orders as a proxy of host/prey diversity for natural enemies (parasitoids and predators). To counterbalance the fact that the same plant species may be over-represented in the literature, we defined three categories: i) plant species related to two or more arthropod orders, ii) plant species related to one arthropod order only, iii) plant species with no data available. These three categories were defined for both floral visitors and phytophagous according to the literature (Supplementary Material Reference List 2S). We gave higher relative importance to the capability of plant species of attracting floral visitors than natural enemies due to biological and technical reasons. Pollinators are key organisms since most plant populations depend on them for not being at risk (Ollerton et al., 2011; Rodger et al., 2021). Furthermore, and whereas immersed in a global context of pollinator decline (Potts et al., 2010), green roofs appear as a promising strategy to promote their food sources in cities (e.g., Wang et al., 2017; Kratschmer et al., 2018). In addition, and although natural enemies play an important role in helping plants to control pests, we assumed that the food resources for phytophagous will be not as scarce as for pollinators given that the former depend on leaves, a resource that is less transient than flowers, and that several groups of herbivores are not detrimentally affected by urbanization (Raupp et al., 2010). Flowers, in addition, may be food resources for both pollinators and natural enemies (Campbell et al., 2012). Lastly, for pollinators, we were able to gather direct information of the resources consumed, whereas for natural enemies a proxy through availability of phytophagous was used, which reinforces the priority we gave to pollinators over natural enemies in the selection criteria.

The Analytic Hierarchy Process (AHP) was then used to define the criteria weights, by means of a pairwise comparison matrix of the relative importance (e.g., the importance of habitat affinity relative to colonization potential, occurrence and performance). From this process, we obtained the criteria weights. Details on the calculus of the criteria weights are given in Appendix A, which defined the following relative order from the most to the least important criteria we have: Habitat affinity > Colonization potential > Performance > Occurrence in the first round, and Tolerance to green roof conditions> Floral visitors attraction potential > Natural enemies attraction potential in the second round (Figure 2). We further ranked the native and exotic species in the final list by PROMETHEE. We performed the ranking procedures for natives and exotics separately. Further details on the PROMETHEE procedure are given in the Statistical analyses section.

### 2.3 Selection of ranked plant species to be tested on roofs

In order to obtain the best-ranked species from both natives and exotics, we introduced an average mark that indicates the position that a plant species with average trait values for all the ranking criteria would have. Therefore, only species above the mark were suitable to be selected. Afterwards, we chose the species to be established in the experimental green roofs from that pool. We selected six native and six exotic plant species from each list giving priority to co-generic, co-familiar plant species, or species with similar traits (e.g., succulence) whenever possible to arrive at two similar species’ pools irrespective of their position above the mark.

### 2.4 Experimental green roofs

The experiment was carried out in Córdoba city, Argentina, from August 2018 to March 2020. A call for volunteers to participate in the experiment was performed from September to November 2018 through social networks. After interviews with 106 volunteers and visits to the roofs, 30 houses were selected for the experimental setting, based on characteristics of their roofs. Selected roofs had a minimum size of 15m^2^, and a height between 3 to 3.5m. For logistic reasons, the degree of accessibility of roofs and location in the city was also considered, as well as the time availability of the owners. The final selected roofs were distributed all over the city (Supplementary Material Figure S1).

The selected native and exotic species were grown from August to November 2018 at the nursery of the Universidad Nacional de Córdoba (Figure 1). To do so, we first collected propagules (seeds or rhizomes) of each native species from urbanized populations whenever possible, given that urban provenance may contribute to species survival in the city (e.g., Yakub and Tiffin, 2017). Only one native species was obtained from a national ornamental variety available in the market (see Results), as we tested this variety previously for the occurrence of floral visitors (Calviño, unpublished results). Most exotics, in addition, were introduced either as rhizomes or directly obtained as plantings from wholesale nurseries. Most importantly, and despite the introduction method may influence the future plants’ performance (e.g., Ksiazek-Mikenas et al., 2021), we did not find any differences in plants’ success related to their introduction method (Calviño et al., unpublished results).

A modular green roof system (medium-density polyethylene 50 × 50 × 15cm modules) was selected to be installed in each of the 30 selected roofs, during February 2019. The substrate was composed of vermiculite, peat moss and compost (1:1:1). Each species was planted in two modules with an initial cover of 0.16m^2^ per species (N=720 modules in total). Before green roof installation, plants were able to grow in their definitive modules for two months for final rustication. Two blocks of 12 modules each (two modules per species) containing the native or exotic plant assemblages and separated by 2.5m, were installed on each roof. All experimental green roofs were initially watered after establishment, and further, we split the 30 experimental roofs into two groups with 15 roofs each (Figure 1). In one group, the plants were regularly watered and spontaneous species were weeded for one year (hereafter WW treatment). In the second group, plants were left without watering or weeding (noWW treatment) for the same period. One year after installation, we recorded the species’ occurrence in their original modules (N=720) and the total cover in square meters reached at the end of the essay by each species, considering the two modules together (N=320). Plant cover was estimated from digital pictures of the modules taken at 1m height using the ImageJ software (Schneider et al., 2012).

### 2.5 Statistical analyses

All statistical analyses were performed in the R environment (version 3.6.1; R Core Team, 2019). We used PROMETHEE (MCDA package; Meyer et al., 2021) as the outranking method. PROMETHEE I and II functions (Bigaret et al., 2017; Meyer et al., 2021) were used to obtain the partial preorder and the complete order, respectively (Brans et al., 1986), on the basis of the criteria weights defined by AHP (Appendix A).

To test the effect of plant origin and management (WW vs. noWW) on species occurrence and cover we first performed generalized linear mixed-effects models (glmer function from the lme4 package; Bates et al., 2018) with roof as a random term and plant origin, management and their interaction as fixed effects, assuming a binomial and Gaussian distribution of errors for plant occurrence and cover, respectively. However, we further decided not to include the random term into the models, given that it accounted for a variance near zero (9.3 × 10^−5^±0.06). In addition, we compared the effect of management on the cover of each of the 12 species one year after establishment, to assess the species’ performance irrespective of their origin. To do so we performed a generalized linear model with management, plant species and their interaction as predictors assuming Gaussian error distribution for the response variable. In all cases, the significance of predictor variables was determined by deviance tests with α=0.05 for significant effects and 0.05<α<0.09 for marginally significant effects. Predicted values of each of the models were plotted with the *sjPlot* package (Lüdecke, 2021).

## 3. Results and Discussion

### 3.1 Same decision framework, different outcomes: the origin effect

Urban green design faces many challenges given the complex decisions involved in planning (Saaty and De Paola, 2017), especially when the goal is to conserve urban wildlife. So far, the decisions to use native or exotic plant species in green roofs had never considered the plant potential to promote beneficial arthropods using the same selection framework. By combining the habitat template hypothesis with surrogates of plant affinities for arthropods in a multicriteria decision framework, we obtained a ranked set of candidate native and exotic plant species expected to tolerate roof conditions and able to attract floral visitors and natural enemies (Supplementary Material Ranked species lists). After the second outranking process, 29 native and 28 exotic plant species were ranked above the mark (Appendix B). Within natives, all of these 29 species were registered both as growing in rocky and in ruderal habitats. Regarding exotics, only *Sedum mexicanum* was identified as growing in both types of habitats. Most of the exotics (23 of 28) were registered as ruderal, including ornamental species registered as escaped from cultivation (e.g., *Verbena hybrida, Zinnia elegans*). In addition, most of the species (Appendix B) have the potential for attracting floral visitors from the order Hymenoptera (72% of the natives and 84% of the exotics) and phytophagous from the order Hemiptera as prey for natural enemies (72% of the natives and 78% of the exotics).

From the species situated above the mark, we chose a pool of six native and six exotic plant species to experimentally test their performance under two contrasting management conditions (i.e., WW and noWW; Table 1). We showed that irrigation and weeding of spontaneous plants clearly benefited the occurrence and cover of both natives and exotics one year after establishment. Particularly, the effect of management on plant occurrence depended on plant origin (interaction term: *D*=3.87, *P*=0.049). According to our expectations based on the adaptation argument (Butler et al., 2012), natives showed an advantage over exotics under noWW since they were more likely to occur under this treatment (Figure 3). Plant origin also had a marginally significant effect on plant cover (*D=0*.*10, P*=0.09; Figure 4A), with natives having a slightly higher cover than exotics one year after establishment. However, the effect of management, in this case, was independent (*D*=0.02, *P*=0.38) and more pronounced than plant origin since plants exhibited a 2.5-fold increase in cover under WW compared with noWW treatment (*D=5*.*14, P*<0.0001; Figure 4B). Looking at the individual performance of each plant species, the model indicates that there was a significant interaction between species and management (*D*=1.48, *P*<0.001). All plant species surpassed their initial cover after one year of establishment, with seven of them surpassing their original cover only under the WW treatment and three natives and two exotics surpassing their initial cover even under the noWW treatment (Figure 5).

**Table 1.**
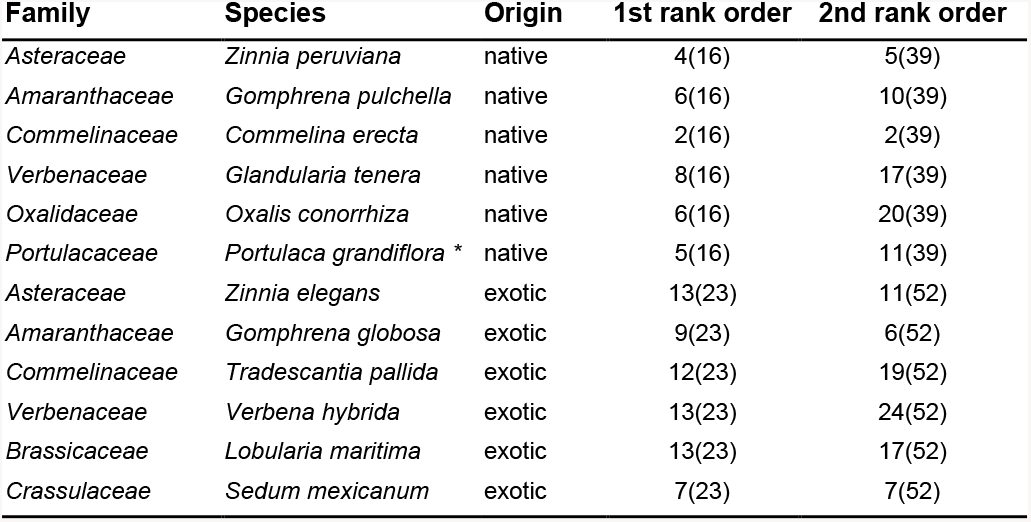
Native and exotic plant species established on the experimental green roofs. First and second rank orders indicate the absolute position after the first and second PROMETHEE analyses, respectively. The total number of ranking categories obtained after each procedure is between brackets (i.e., distinct plant species may arrive at the same rank position). Please, see Methods for further details. * corresponding to *P. grandiflora* ‘INTA’ in our experiment.

**Figure 3.**
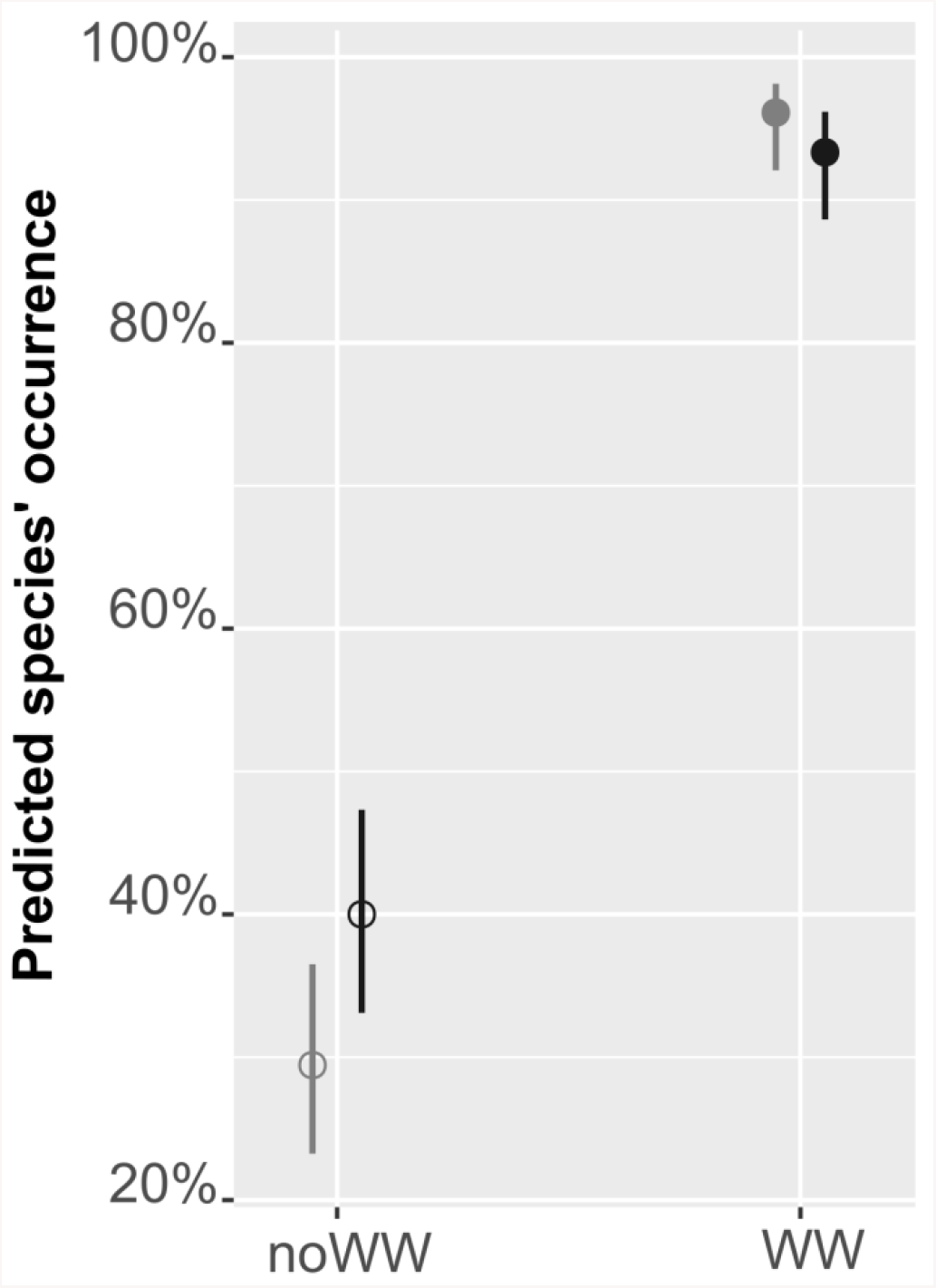
Predicted species occurrences per module in relation to origin (natives in black, exotics in grey), and management (WW= watering and weeding, filled circles, noWW=no watering nor weeding of spontaneous plants, empty circles), one year after establishment in the experimental green roofs.

**Figure 4.**
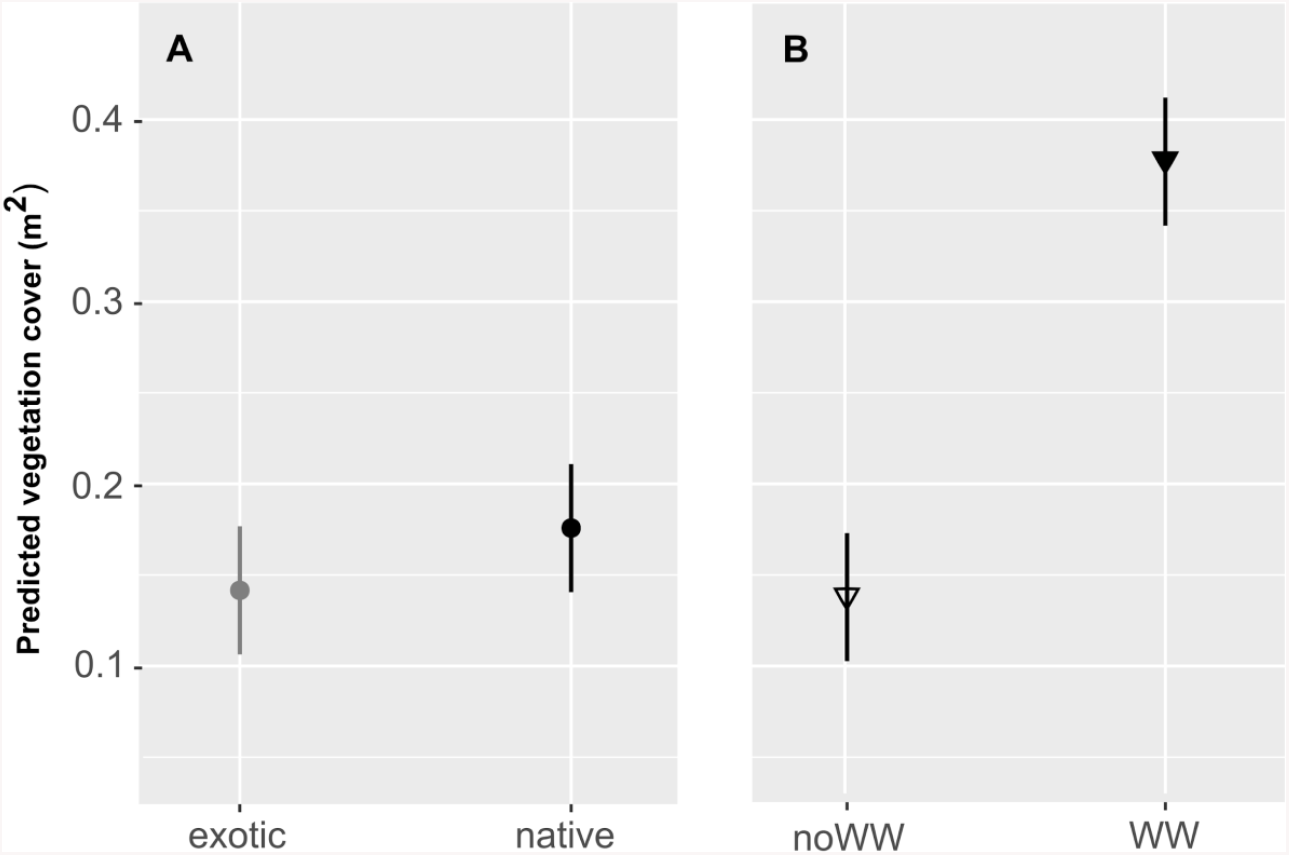
Predicted species cover (m^2^) one year after establishment in the experimental green roofs in relation to A). Plant origin: native (black) and exotic (grey). B) Plant management: under watering and weeding (WW, filled symbol) and with no watering nor weeding of spontaneous plants (noWW, empty symbol).

**Figure 5.**
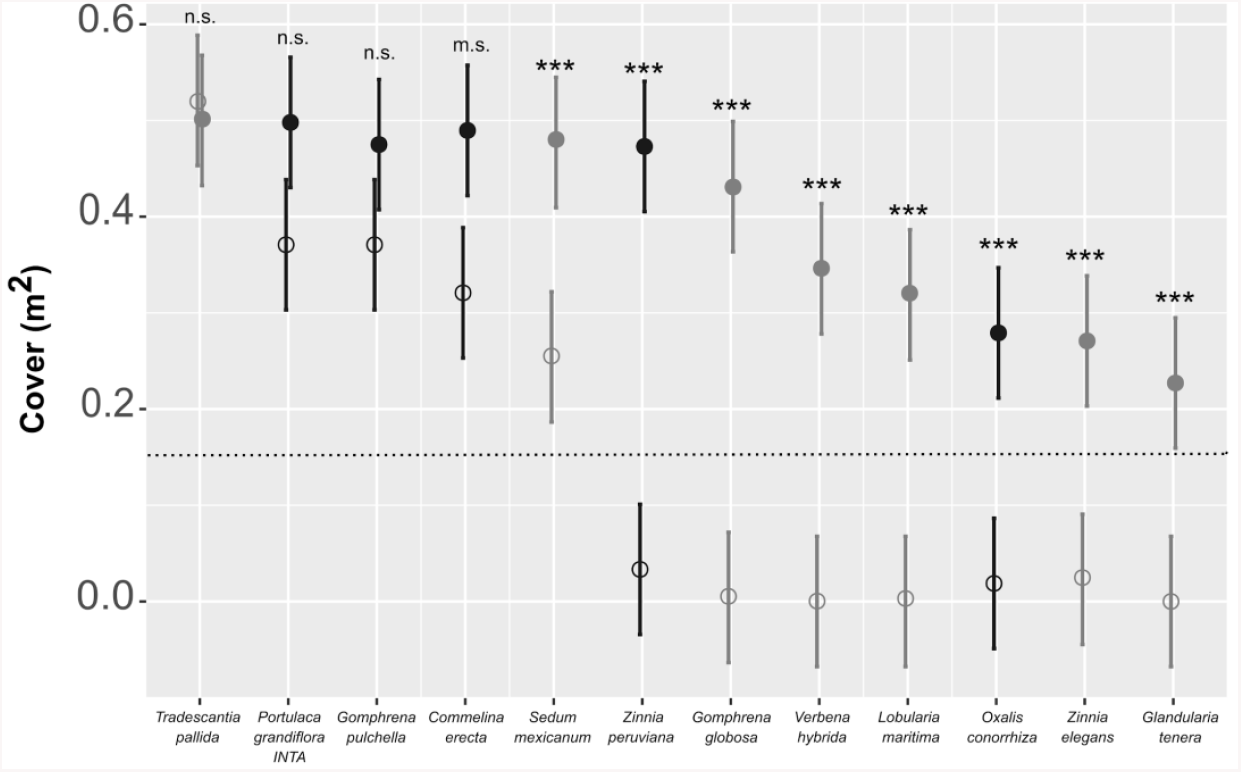
Individual plant species cover (m^2^) one year after the experimental green roof installation. From left to right, species are ordered from their highest to their lowest final cover. In black: roofs with regular watering and weeding, and in grey control roofs (no watering nor weeding during the same period). Circular marks represent native species, and triangles exotic ones. Significant changes in cover within each species given due to management effects were obtained using a post hoc Tuckey test and are denoted by asterisks. \*\**P*=0.01; \*\*\**P*≤0.0001, m.s.=marginally significant, n.s.= non-significant. The horizontal line is the initial plant cover at the establishment (0.16m^2^).

### 3.2 Winners and losers: a false dichotomy?

Under the semiarid climate of Córdoba city (Cwa in the Köppen-Geiger Climatic Classification; Beck et al., 2018) and despite the limited number of species tested along just one year, our results shed light on the importance of choosing natives for future extensive green roofs. In fact, two of the annual native species here evaluated (*P. grandiflora* and *G. pulchella*) were able to reseed after winter even in the absence of irrigation, reaching similar cover levels to those registered by the same species under irrigation and weeding. This result illustrates how some native annuals are capable of reseeding after the dry winter season in the experimental green roofs, just as they do in their natural habitats. Furthermore, and considering that our treatment of management represents two contrasting conditions (regular watering and weeding vs no intervention), we expect that intermediate or even minor irrigation levels may broaden the spectrum of plant species suitable for green roofs. This is likely the case of the native *Z. peruviana* or the exotic *G. globosa*, two annual species that exhibited the greatest differences in cover between the two treatments (i.e., higher reseeding capacity only under irrigation and weeding). *Zinnia peruviana* and *G. globosa* had a great potential to reseed after winter under WW, with a three and a 2.6-fold increase from their initial cover, respectively. These results are in agreement with Zhang et al. (2021), who found self-sowing is a good surrogate of plant resilience in green roofs. On the other hand, it is interesting that some of the native and exotic plant species that did not perform well, especially in the absence of irrigation, are perennials (Appendix B). Although we did not include plant life span in our decision framework, it seems that it is a key trait to design low maintenance extensive green roofs in Córdoba city. However, future studies that consider life span in an integrative framework are necessary to test this idea.

Regarding the ability to succeed or not in the roofs, other aspects deserve to be explored. First, our results coincide with previous findings that highlight the importance of succulence in urban environments given their well-known high survival and recovery from drought (reviewed in Lundholm and Walker, 2018; but see Guo et al., 2021). One native (*P. grandiflora*) and one exotic (*S. mexicanum*) succulent plant species were among the plants with the highest cover values. This may also be true for the exotic *T. pallida* with a rather low succulence degree, but it should be taken with caution given its recent spread in the city (Calviño, obs. pers.). Second, it is important to mention that new hybrid varieties were recently obtained for the native *Glandularia* spp. (Suárez, 2020) just after this experiment was established, and it is expected that these varieties would exhibit better performance than the wild relative *G. tenera* here tested (e.g., Henson et al. 2006; for *G. tenuisecta* x *G. tenera* hybrid). Nevertheless, selection with an ornamental purpose only could be detrimental for some plant-insect interactions (e.g., Mach and Potter, 2018) and tests on the new hybrids should be really helpful in this regard.

Overall, and given that all species surpassed their original cover, our results are in agreement with those of Yee et al. (2021) in that relevant plant species for green roof ecosystems should not necessarily be the most abundant. Most importantly, a ranking process like the one used here may be a helpful tool to overcome traditional selection criteria and include plant species that tolerate the roofs and are also able to attract urban wildlife.

### 3.3 Integrating green roof design with biodiversity conservation goals

As Sikorski et al. (2021) have shown for informal urban green spaces, minimal interventions could guarantee several ecosystem benefits and even favour local biodiversity, a set of outcomes usually associated with more costly management and design strategies. In this regard, and considering that two annual species here tested were able to reseed without regular watering and that adding annuals has positive effects on green roof arthropods (e.g., Salman and Blaustein, 2018), we sustain that the relative success of a given species should be considered in a broader sense, integrating the potential of a given species to foster urban biodiversity. Urban environments are usually characterized by their restricted value for animals (Apfelbeck et al., 2019), especially for insects (Egerer and Buchholz, 2021; Fenoglio et al., 2021). Therefore, the idea that only minimal interventions are needed to favour arthropods is especially attractive for their conservation in cities. In this regard, we do not ignore that most plant species studied here have some negative social perceptions. Although they may establish crucial trophic relationships with different insect groups, most of the best ranked native plant species we obtained are considered weeds in our country, particularly in agricultural habitats. For instance, *G. pulchella* is well recognized as a herbicide-tolerant weed (Calderón, 2013) but presents a high butterfly attractiveness. *Commelina erecta*, another successful species in our study, is able to sustain a high diversity of floral visitors and natural enemies according to the literature (Fenoglio et al., 2010; Faden, 1992) but is also a glyphosate-tolerant undesirable weed (e.g., Gullino et al., 2016). In landscape design, plants with both a “natural affinity and a strong visual relationship” are considered as plant signatures (Robinson, 1993). Often found at rural roadsides, both G. pulchella and C. erecta may appear associated with Z. peruviana and P. grandiflora in ruderal habitats (Calviño, pers. obs.). This confirms the “close identity between plants and place”, a key aspect of the plants’ signature concept of Robinson (1993), that could be useful to reframe urban wildlife (Egerer and Buchholz, 2021). Ruderal, formerly “weeds”, can be reframed as “pollinator attractive plants’’ and “beneficial insectary plants” (i.e. plants supporting alternate hosts for predators and parasitoids according to Atsatt & O’Dowd, 1976) to be included in green roofs design. This reframing may provide beneficial signatures not only to urban arthropod wildlife but also to broaden the spectrum of native species typically chosen to be established in green roofs.

## 4. Concluding remarks

Our work gives tools for green roof design that help to select native and exotic plant species by using, for the first time, a consistent multicriteria decision-making approach combining traditional with novel criteria which mirror the potential of plants for promoting arthropod biodiversity. In addition, by experimentally evaluating 12 candidate species obtained after applying the MCDA, we showed that native species performed better than exotics according to what is expected under the “adaptation argument” (Butler et al., 2012). These results constitute new evidence for a South American city where green roof technology and, especially, the selection and use of native vegetation are taking their first steps (but see Jaramillo Pazmino, 2016; Cáceres et al., 2018). In addition, given that the robustness of the MCDA procedures was tested in only 5% of the studies reviewed in conservation (Adem Esmail and Geneletti, 2018), our results contribute to addressing the utility of this approach to select plants for extensive green roof design aimed to favour urban biodiversity. Although we used a limited number of plant species in our essay and future studies are necessary to test the real power of arthropod attraction by plants depending on their origin, our study goes one step forward on the current methods used for plant selection in green roof design and sheds light into the importance of choosing natives for future extensive green roofs. Considering green roofs are one of the possible solutions to ameliorate the negative effects of urban habitat loss on arthropod diversity (Fenoglio et al., 2021), the development of an integrative multicriteria decision framework that takes into account the potential of both native and exotic plant species for promoting beneficial arthropods would give a new twist in plant selection processes for green roofs.

## Acknowledgements

We are very grateful to all the volunteers that offered their houses to install the experimental green roofs. We thank Gustavo Bertone, Agostina Bordunale, Martin Videla, Mia Martin, and Mateo Barrera for their help with the fieldwork. We also thank Ezequiel González for his very helpful comments on the early versions of the manuscript. This work was supported by funds from the National Geographic Society (NGS-383R-18 grant awarded to MSF). AAC, AS, EE, HMB, MSF, and MLM are scientific researchers at the Consejo Nacional de Investigaciones Científicas y Técnicas (CONICET-Universidad Nacional de Córdoba); DF is a doctoral fellow of CONICET at the Universidad Nacional de Córdoba and JT is a postdoctoral fellow of the Royal Society of London.

## Appendix A

**Table A1.** Numerical categorization of the criteria used in the AHP model of Figure 1.

| **Selection and ranking criteria** | **Description** |
| --- | --- |
| Habitat affinity | Represents the theoretical similarity of rooftops with natural habitats, and it was obtained by summing up values assigned to three different habitat templates: ruderal (value=1 if classified as ruderal, either equal to 0); rocky (value=1 if classified as stonecrop or rocky habitats, either equal to 0); another habitat (value=0.5 if any other habitat was registered like grassy, mountainous, coastlands, etc.). |
| Occurrence | The number of studies in which the species was registered in a green roof over the maximum number observed over all plant species |
| Colonization potential | Whether the species was registered as spontaneous colonizer (value=1) or not (value=0). |
| Performance | Whether the data available support either 50% of cover, germination rate or roofs' frequency in a given species: value= 1, whether the data support less than 50% or there are no data available: value=0.5 (*). |
| Floral visitors' attraction potential | Plant species related to one insect order= 1<br>Plant species related to two or more insect orders=2<br>Plant species with no data available=0 |
| Natural enemies' attraction potential | Plant species related to one insect order= 1<br>Plant species related to two or more insect orders=2<br>Plant species with no data available=0 |

(\*) No data was not considered a "zero" value given that we started the selection procedure with candidate species already registered as growing in green roofs.

### Paired comparison by Analytic hierarchy process and criteria weight definition

The Analytic hierarchy process (AHP) was used to define the relative criteria weights used further in the PROMETHEE ranking method. Here we describe the method of Saaty (2008) we used to obtain the weights of the criteria illustrated in Figure 1.

The weights of the criteria used in PROMETHEE were equal to the priority vector resulting from the pairwise comparison of the criteria, the main procedure of the AHP. Originally, the AHP was used to express different judgements in the form of comparisons, a method especially useful to define decision priorities in situations involving many people (e.g., Saaty, 2004). Thus, the first step of the method consists of assigning to each pair, the importance of one criterium relative to the importance of the other criterium, according to a scale. The original Saaty’s scale was divided into nine “intensities of importance” with equal importance =1, and higher numbers representing stronger importance, however, other possibilities may be useful depending on the parameter’s variability (Saaty, 1990, 2004). Here, 4 × 4 and 3 × 3 pairwise comparison matrices were used for the first and second rounds, respectively (Figure 1). A shorter version of the 9-scale of Saaty was used with only four categories. For instance, habitat affinity was four times more important than occurrence, and the reciprocal means that occurrence was ¼ times more important relative to habitat affinity (Table A2). The priority vector represents the relative importance of each criterion in the whole matrix (i.e., criteria weights), and it is equal to the row averages of the normalized matrix (Saaty, 1990). Accordingly, and from the most to the least important criteria we have: Habitat affinity> Colonization potential> Performance> Occurrence, for the first round, and Tolerance to green roof conditions for the second round> Floral visitors’ attraction potential >Natural enemies’ attraction potential, for the second round (Figure 2).

**Table A2.** Matrix for the pairwise comparison used to estimate the criteria weight in the first decision round.

|  | Occurrence | Colonization potential | Performance | Habitat affinity | Priority vector |
| --- | --- | --- | --- | --- | --- |
| Occurrence | 1 | 0.33 | 0.5 | 0.25 | 0.097 |
| Colonization potential | 3 | 1 | 1.5 | 0.33 | 0.245 |
| Performance | 2 | 0.66 | 1 | 0.66 | 0.213 |
| Habitat affinity | 4 | 3 | 1.5 | 1 | 0.444 |

**Table A3.** Matrix for the pairwise comparison used to estimate the criteria weight in the second decision round.

|  | Tolerance to green roof conditions | Floral visitors' attraction potential | Natural enemies' attraction potential | Priority vector |
| --- | --- | --- | --- | --- |
| Tolerance to green roof conditions | 1 | 3 | 4 | 0.623 |
| Floral visitors' attraction potential | 0.33 | 1 | 2 | 0.239 |
| Natural enemies' attraction potential | 0.25 | 0.5 | 1 | 0.137 |

It is further important to note that judgement’ inconsistencies may arrive from the pairwise comparison matrix and, as Saaty (2004) has pointed out, “consistency is essential in human thinking because it enables us to order the world according to dominance”. To overcome a potential inconsistency problem, we obtained the consistency ratio (CR) for each matrix. The CR was obtained as the ratio between the consistency index and a randomly generated index (Saaty, 1980). CR were 0.045 and 0.016 for the first and second pairwise comparisons, respectively. Given that CR was less than 0.1 for both matrices, inconsistencies are less than 10% and our judgments are therefore confident (Saaty, 1980).

## Appendix B.

Candidate plant species for green roofs obtained after applying the integrative multicriteria decision framework. Rank order indicates the absolute position of the species after the second PROMETHEE decision round above the mark (i.e., average imaginary species). The taxonomic order of floral visitors and phytophagous were assigned according to the references in Table S1. The final species selected to be established in the experimental green roofs are in bold. RO= Rocky outcrop habitats, RU= Ruderal, O= Other (sandy, grasslands, mountainous, etc.). Na= No data.

| Rank order | Family | Name | Origin | Lifespan | Habitat analogs | Floral visitors | Phytophagous |
| --- | --- | --- | --- | --- | --- | --- | --- |
| 1 | Asteraceae | *Bidens pilosa* | native | annual | RU, RO | Coleoptera, Diptera, Hymenoptera, Lepidoptera | Coleoptera, Diptera, Hemiptera, Thysanoptera |
| **2** | **Commelinaceae** | ***Commelina erecta*** | **native** | **perennial** | **RU, RO** | **Coleoptera, Diptera, Hymenoptera, Lepidoptera** | **Coleoptera, Diptera, Hemiptera** |
| 3 | Convolvulaceae | *Ipomoea purpurea* | native | annual | RU, RO | Diptera, Hymenoptera | Coleoptera, Diptera, Hemiptera |
| 4 | Acanthaceae | *Dicliptera squarrosa* | native | perennial | RU, RO | Diptera, Hymenoptera, Lepidoptera | Diptera, Hemiptera, Lepidoptera |
| 4 | Asteraceae | *Bidens subalternans* | native | annual | RU, RO | Diptera, Hymenoptera, Lepidoptera | Coleoptera, Hemiptera, Lepidoptera |
| 4 | Asteraceae | *Conyza bonariensis* | native | annual | RU, RO | Coleoptera, Diptera, Hymenoptera, Lepidoptera | Coleoptera, Diptera, Hemiptera, Hymenoptera, Lepidoptera, Orthoptera |
| 4 | Malvaceae | *Malvastrum coromandelianum* | native | perennial | RU, RO | Hymenoptera, Lepidoptera | Coleoptera, Diptera, Hemiptera, Lepidoptera, Thysanoptera |
| 4 | Malvaceae | *Sida rhombifolia* | native | perennial | RU, RO | Coleoptera, Hymenoptera, Lepidoptera | Coleoptera, Diptera, Hemiptera, Lepidoptera, Thysanoptera |
| 4 | Malvaceae | *Sida spinosa* | native | perennial | RU, RO | Hymenoptera, Lepidoptera | Coleoptera, Diptera, Hemiptera, Lepidoptera, Thysanoptera |
| 5 | Asteraceae | *Conyza sumatrensis* | native | annual | RU, RO | Diptera, Hymenoptera, Lepidoptera | Diptera, Hemiptera |
| **5** | **Asteraceae** | ***Zinnia peruviana*** | **native** | **annual** | **RU, RO** | **Hymenoptera, Lepidoptera** | **Diptera, Hemiptera** |
| 6 | Lythraceae | *Heimia salicifolia* | native | perennial | RU, RO | Hymenoptera | Hemiptera, Hymenoptera, Lepidoptera |
| 7 | Convolvulaceae | *Ipomoea nil* | native | annual | RU, RO | Coleoptera, Diptera, Hymenoptera | Coleoptera, Hemiptera |
| 8 | Convolvulaceae | *Ipomoea cairica* | native | perennial | RU, RO | Coleoptera, Diptera, Hymenoptera | Coleoptera, Diptera, Hemiptera, Lepidoptera, Thysanoptera |
| 9 | Asteraceae | *Parthenium hysterophorus* | native | annual | RU, RO | Na | Coleoptera, Diptera, Hemiptera |
| 9 | Poaceae | *Paspalum dilatatum* | native | perennial | RU, RO | Na | Coleoptera, Hymenoptera, Lepidoptera, Orthoptera, Thysanoptera |
| 10 | **Amaranthaceae** | ***Gomphrena pulchella*** | **native** | **annual** | **RU, RO** | **Hymenoptera, Lepidoptera** | **Hemiptera** |
| 11 | **Portulacaceae** | ***Portulaca grandiflora*** | **native** | **annual** | **RU, RO** | **Hymenoptera** | **Hemiptera** |
| 12 | Convolvulaceae | *Dichondra sericea* | native | perennial | RU, RO | Na | Hymenoptera, Lepidoptera |
| 12 | Poacea | *Eragrostis pastoensis* | native | perennial | RU, RO | Na | Hemiptera, Thysanoptera |
| 13 | Asteraceae | *Schkuhria pinnata* | native | annual | RU, RO | Hymenoptera | Thysanoptera |
| 13 | Solanaceae | *Nierembergia linariaefolia* | native | perennial | RU, RO | Hymenoptera | Thysanoptera |
| 14 | Asteraceae | *Tagetes minuta* | native | annual | RU, RO | Na | Hemiptera |
| 15 | Cyperaceae | *Cyperus rotundus* | native | perennial | RU | Hymenoptera | Coleoptera, Hemiptera, Lepidoptera, Orthoptera |
| 16 | Asteraceae | *Galinsoga parviflora* | native | annual | RU | Coleoptera, Diptera, Hymenoptera, Lepidoptera | Hemiptera, Thysanoptera |
| 17 | **Verbenaceae** | ***Glandularia tenera*** | **native** | **perennial** | **RU, RO** | **Hymenoptera, Lepidoptera** | **Lepidoptera** |
| 18 | Cyperaceae | *Cyperus aggregatus* | native | perennial | RU, RO | Na | Orthoptera |
| 19 | Poaceae | *Setaria parviflora* | native | perennial | RU, RO | Na | Hemiptera |
| 20 | **Oxalidaceae** | ***Oxalis conorrhiza*** | **native** | **perennial** | **RU, RO** | **Hymenoptera** | **Na** |
| 1 | Asteraceae | *Taraxacum officinale* | exotic | perennial | RU | Coleoptera, Diptera, Hymenoptera, Lepidoptera | Coleoptera, Diptera, Hemiptera, Hymenoptera, Lepidoptera, Orthoptera, Thysanoptera |
| 2 | Chenopodiaceae | *Chenopodium album* | exotic | annual | RU | Coleoptera, Diptera, Hymenoptera | Coleoptera, Diptera, Hemiptera, Lepidoptera, Orthoptera, Thysanoptera |
| 3 | Aizoaceae | *Aptenia cordifolia* | exotic | perennial | RU, O | Hymenoptera, Lepidoptera | Hemiptera |
| 4 | Poacea | *Digitaria sanguinalis* | exotic | annual | RU | Hymenoptera | Coleoptera, Hemiptera, Lepidoptera, Orthoptera |
| 5 | Amaryllidaceae | *Allium schoenoprasum* | exotic | perennial | RO, O | Hymenoptera | Hemiptera |
| 6 | **Amaranthaceae** | ***Gomphrena globosa*** | exotic | annual | RU, O | **Hymenoptera, Lepidoptera** | **Hemiptera** |
| 7 | **Crassulaceae** | ***Sedum mexicanum*** | exotic | perennial | RU, RO | **Hymenoptera** | **Hemiptera** |
| 8 | Asteraceae | *Hypochaeris radicata* | exotic | perennial | RU | Diptera, Hymenoptera | Hemiptera, Thysanoptera |
| 8 | Asteraceae | *Matricaria chamomilla* | exotic | annual | RU | Hymenoptera, Lepidoptera | Hemiptera, Thysanoptera |
| 9 | Xanthorrhoeaceae | *Bulbine frutescens* | exotic | perennial | RO, O | Hymenoptera | Na |
| 10 | Crassulaceae | *Bryophyllum daigremontianum* | exotic | perennial | RU, O | Na | Hemiptera |
| 11 | **Asteraceae** | ***Zinnia elegans*** | exotic | annual | RU, O | **Hymenoptera, Lepidoptera** | **Diptera, Hemiptera** |
| 12 | Portulacaceae | *Portulaca oleracea* | exotic | annual | RU | Hymenoptera | Coleoptera, Hemiptera, Hymenoptera, Lepidoptera, Thysanoptera |
| 13 | Crassulaceae | *Sedum acre* | exotic | perennial | RO | Hymenoptera | Na |
| 14 | Plantaginaceae | *Plantago major* | exotic | perennial | RU | Na | Hemiptera |
| 14 | Oxalidaceae | *Oxalis corniculata* | exotic | perennial | RU | Na | Coleoptera |
| 15 | Fabaceae | *Trifolium repens* | exotic | perennial | RU | Hymenoptera, Lepidoptera | Coleoptera, Hemiptera, Hymenoptera, Lepidoptera, Thysanoptera |
| 16 | Lamiaceae | *Leonurus japonicus* | exotic | biennial | RU | Hymenoptera, Lepidoptera | Hemiptera, Thysanoptera |
| 17 | **Brassicaceae** | ***Lobularia maritima*** | exotic | perennial | RO | **Diptera, Hymenoptera** | **Hemiptera** |
| 18 | Papaveraceae | *Papaver rhoeas* | exotic | annual | RU | Hymenoptera | Diptera |
| 19 | **Commelinaceae** | ***Tradescantia pallida*** | exotic | perennial | RU | **Hymenoptera** | **Hemiptera** |
| 20 | Asteraceae | *Leucanthemum vulgare* | exotic | perennial | RU | Diptera, Hymenoptera | Coleoptera, Hemiptera, Lepidoptera |
| 21 | Crassulaceae | *Sedum album* | exotic | perennial | RO | Hymenoptera | Na |
| 22 | Poaceae | *Chloris gayana* | exotic | perennial | RU | Hymenoptera | Hemiptera, Thysanoptera |
| 23 | Poacea | *Echinochloa colona* | exotic | annual | RU | Na | Coleoptera, Hemiptera |
| 24 | Verbenaceae | *Verbena hybrida* | exotic | perennial | RU | Diptera, Hymenoptera, Lepidoptera | Diptera, Hemiptera |
| 24 | Amaryllidaceae | *Allium ampeloprasum* | exotic | perennial | RU | Diptera, Hymenoptera | Hemiptera, Thysanoptera |
| 24 | Plantaginaceae | *Veronica persica* | exotic | perennial | RU | Diptera, Hymenoptera | Hemiptera, Thysanoptera |

## Notes

### Competing Interest Statement

Maria Silvina Fenoglio reports financial support was provided by National Geographic Society.

## References

Adem Esmail, B., Geneletti, D., 2018. Multi-criteria decision analysis for nature conservation: A review of 20 years of applications. Methods Ecol. Evol. 9, 42–53. https://doi.org/10.1111/2041-210X.12899

Apfelbeck, B., Jakoby, C., Hanusch, M., Steffani, E.B., Hauck, T.E., Weisser, W.W., 2019. A conceptual framework for choosing target species for wildlife-inclusive urban design. Sustainability. 11, 6972. https://doi.org/10.3390/su11246972

Asgarzadeh, M., Vahdati, K., Lotfi, M., Arab, M., Babaei, A., Naderi, F., Pir Soufie, M., Rouhani, G., 2014. Plant selection method for urban landscapes of semi-arid cities (a case study of Tehran). Urban For. Urban Green. 13, 450–458. https://doi.org/10.1016/j.ufug.2014.04.006

Atsatt, P. R., O’Dowd, D. J. (1976). Plant defense guilds. Science. 193, 24–29. https://doi.org/10.1126/science.193.4247.24

Beck, H.E., Zimmermann, N.E., McVicar, T.R., Vergopolan, N., Berg, A., Wood, E.F., 2018. Present and future Köppen-Geiger climate classification maps at 1-km resolution. Sci. Data. 5, 1–12. https://doi.org/10.1038/sdata.2018.214

Berthon, K., Thomas, F., Bekessy, S., 2021. The role of ‘nativeness’ in urban greening to support animal biodiversity. Landsc. Urban. Plan. 205, 103959. https://doi.org/10.1016/j.landurbplan.2020.103959

Bigaret, S., Hodgett, R.E., Meyer, P., Mironova, T., Olteanu, A.L., 2017. Supporting the multi-criteria decision aiding process: R and the MCDA package. EJDP. 5, 169–194. https://doi.org/10.1007/s40070-017-0064-1

Brans, J.P., Vincke, P., Mareschal, B., 1986. How to select and how to rank projects: The PROMETHEE method. Eur. J. Oper. Res. 24, 228–238. https://doi.org/10.1016/0377-2217(86)90044-5

Butler, C., Orians, C.M., 2011. Sedum cools soil and can improve neighboring plant performance during water deficit on a green roof. Ecol. Eng. 37, 1796–1803. https://doi.org/10.1016/j.ecoleng.2011.06.025

Butler, C., Butler, E., Orians, C. M. 2012. Native plant enthusiasm reaches new heights: Perceptions, evidence, and the future of green roofs. Urban For. Urban Green. 11, 1–10. https://doi.org/10.1016/j.ufug.2011.11.002

Cáceres, N., Imhof, L., Suárez, M., Hick, E.C., Galetto, L., 2018. Assessing native germplasm for extensive green roof systems of semiarid regions. Ornam. Hortic. 24, 466–476. http://dx.doi.org/10.14295/oh.v24i4.1225

Calderón, M., 2013. Evaluación de la respuesta de malezas a la aplicación de glifosato en un cultivo de soja (Glycine max) en Victoria, Entre Ríos. Bachelor’s thesis, Universidad Católica Argentina, Paraná. https://repositorio.uca.edu.ar/handle/123456789/375

Campbell, A. J., Biesmeijer, J. C., Varma, V., Wäckers, F. L. (2012). Realising multiple ecosystem services based on the response of three beneficial insect groups to floral traits and trait diversity. Basic and Applied Ecology, 13, 363–370. https://doi.org/10.1016/j.baae.2012.04.003

Cascone, S., 2019. Green roof design: State of the art on technology and materials. Sustainability. 11, 3020. https://doi.org/10.3390/su11113020

Cook-Patton, S.C., 2015. Plant biodiversity on green roofs, in: Sutton, R. (Ed.), Green roof ecosystems Springer, Cham, pp. 193–209. https://doi.org/10.1007/978-3-319-14983-7_8

Dabija, A.M., 2019. Living Envelopes for Buildings–A Historic Parallel. IOP Conf. Ser.: Mater. Sci. Eng. 471, 112014. https://doi.org/10.1088/1757-899X/471/11/112014

Dimitri, M.J., Parodi, L.R., 1977. Enciclopedia Argentina de agricultura y jardinería (No. 630). Acme, Buenos Aires.

Egerer, M., Buchholz, S., 2021. Reframing urban “wildlife” to promote inclusive conservation science and practice. Biodivers. Conserv. 30, 2255–2266. https://doi.org/10.1007/s10531-021-02182-y

Fabián, D., González, E., Domínguez, M.V.S., Salvo, A., Fenoglio, M.S., 2021. Towards the design of biodiverse green roofs in Argentina: Assessing key elements for different functional groups of arthropods. Urban For. Urban Green. 61, 127107. https://doi.org/10.1016/j.ufug.2021.127107

Faden, R.B., 1992. Floral Attraction and Floral Hairs in the Commelinaceae. Ann. Missouri Bot. Gard. 79, 46–52. https://doi.org/10.2307/2399808

Fenoglio, M.S., Salvo, A., Videla, M., Valladares, G., 2010. Plant patch structure modifies parasitoid assemblage richness of a specialist herbivore. Ecol. Entomo. 35, 594–601. https://doi.org/10.1111/j.1365-2311.2010.01218.x

Fenoglio, M.S., Calviño, A., González, E., Salvo, A., Videla, M., 2021. Urbanisation drivers and underlying mechanisms of terrestrial insect diversity loss in cities. Ecol. Entomo. 46, 757–771. https://doi.org/10.1111/een.13041

Guarino, R., Andreucci, M.B., Leone, M., Bretzel, F., Pasta, S., Catalano, C., 2021. Urban Services to Ecosystems: An Introduction, in: Catalano, C., Andreucci, M.B., Guarino, R., Bretzel, F., Leone, M., Pasta, S. (Eds.), Urban Services to Ecosystems. Future City, vol 17. Springer, Cham, pp. 1–10. https://doi.org/10.1007/978-3-030-75929-2_1

Gullino, C.A., Gomez, P.A., Lorenzatti, L., 2016. Evaluación del nivel de resistencia de un biotipo de Chloris virgata Sw de la región norte de la Provincia de Córdoba al herbicida glifosato. Bachelor’s thesis, Universidad Nacional de Córdoba, Córdoba. http://hdl.handle.net/11086/4364

Guo, B., Arndt, S., Miller, R., Lu, N., Farrell, C., 2021. Are succulence or trait combinations related to plant survival on hot and dry green roofs?. Urban For. Urban Green., 64, 127248. https://doi.org/10.1016/j.ufug.2021.127248

Heim, A., Xie, G., Lundholm, J., 2021. Functional and Phylogenetic Characteristics of Vegetation: Effects on Constructed Green Infrastructure, in: Catalano, C., Andreucci, M.B., Guarino, R., Bretzel, F., Leone, M., Pasta, S. (Eds), Urban Services to Ecosystems. Future City, vol 17. Springer, Cham, pp. 61–83. https://doi.org/10.1007/978-3-030-75929-2_4

Henson, D. Y., Newman, S. E., Hartley, D. E. (2006). Performance of selected herbaceous annual ornamentals grown at decreasing levels of irrigation. HortScience. 41, 1481–1486. https://doi.org/10.21273/HORTSCI.41.6.1481

Hurrell, J.A., Bazzano, D.H., Delucchi, D.G., 2006. Biota Rioplatense XII Dicotiledóneas herbáceas I, 1st edition.. L.O.L.A. - Literature of Latin America, Buenos Aires.

Hurrell, J A., Bazzano, D.H., Delucchi, D.G., 2007. Biota Rioplatense XII Dicotiledóneas herbáceas II, 1st edition. L.O.L.A. - Literature of Latin America, Buenos Aires.

Hurrell, J. A., Delucchi, G., Correa, M. N., Sánchez, M. I., Roitman, G., Buet Costantino, F., … Tur, N. M. 2009. Flora Rioplatense. Sistemática, ecología y etnobotánica de las plantas vasculares rioplatenses. Parte 3. Monocotiledóneas. 1st edition. L.O.L.A. - Literature of Latin America, Buenos Aires.

Hurrell, J.A., Bayón, N.D., Delucchi, G., 2017. Plantas cultivadas de la Argentina, 1st edition. Hemisferio Sur ediciones, Buenos Aires.

Jaramillo Pazmino, M.L., 2016. Plant selection for green roofs in Quito, Ecuador. Master Thesis of Environment. University of Melbourne, Melbourne. http://hdl.handle.net/11343/119536

Kalarus, K., Halecki, W., Skalski, T., 2019. Both semi-natural and ruderal habitats matter for supporting insect functional diversity in an abandoned quarry in the city of Kraków (S Poland). Urban Ecosyst. 22, 943–953. https://doi.org/10.1007/s11252-019-00869-3

Kiehl, K., Jeschke, D., Schröder, R. 2021. Roof Greening with Native Plant Species of Dry Sandy Grasslands in Northwestern Germany, in: Catalano, C., Andreucci, M.B., Guarino, R., Bretzel, F., Leone, M., Pasta, S. (Eds.), Urban Services to Ecosystems. Future City, vol17. Springer, Cham. https://doi.org/10.1007/978-3-030-75929-2_6

Knapp, S., Schmauck, S., Zehnsdorf, A., 2019. Biodiversity impact of green roofs and constructed wetlands as progressive eco-technologies in urban areas. Sustainability,11, 5846. https://doi.org/10.3390/su11205846

Kratschmer, S., Kriechbaum, M., Pachinger, B., 2018. Buzzing on top: Linking wild bee diversity, abundance and traits with green roof qualities. Urban Ecosyst. 21, 429–446. https://doi.org/10.1007/s11252-017-0726-6

Ksiazek-Mikenas, K., Chaudhary, V.B., Larkin, D.J., Skogen, K.A., 2021. A habitat analog approach establishes native plant communities on green roofs. Ecosphere. 12, e03754. https://doi.org/10.1002/ecs2.3754

Lüdecke, D., 2021. sjPlot: Data Visualization for Statistics in Social Science. R package (version 2.8.8). https://CRAN.R-project.org/package=sjPlot

Lundholm, J.T., 2006. Green roofs and facades: a habitat template approach. Urban habitats. 4, 87–101.

Lundholm, J.T., Walker, E.A., 2018. Evaluating the habitat-template approach applied to green roofs. Urban Naturalist. 1, 39–51.

Mach, B.M., Potter, D.A., 2018. Quantifying bee assemblages and attractiveness of flowering woody landscape plants for urban pollinator conservation. PLoS One. 13, e0208428. https://doi.org/10.1371/journal.pone.0208428

MacIvor, J.S., Ksiazek, K., 2015. Invertebrates on green roofs, in: Sutton, R. (Ed.), Green Roof Ecosystems. Ecological Studies (Analysis and Synthesis), vol 223. Springer, Cham, pp. 333–355. https://doi.org/10.1007/978-3-319-14983-7_14

Marttunen, M., Lienert, J., Belton, V., 2017. Structuring problems for Multi-Criteria Decision Analysis in practice: A literature review of method combinations. Eur. J. Oper. Res. 263, 1–17. https://doi.org/10.1016/j.ejor.2017.04.041

Mata, L., Andersen, A.N., Morán-Ordóñez, A., Hahs, A.K., Backstrom, A., Ives, C.D., Bickel, D., Duncan, D., Palma, E., Thomas, F., Cranney, K., Walker, K., Shears, I., Semeraro, L., Malipatil, M., Moir, M.L., Plein, M., Porch, N., Vesk, P.A., Smith, T.R., Lynch, Y., 2021. Indigenous plants promote insect biodiversity in urban greenspaces. Ecol. Appl. 31, e02309. https://doi.org/10.1002/eap.2309

Meyer, P., Bigaret, S., Richard Hodgett, R., Olteanu, A.L., 2021. MCDA: Support for the Multicriteria Decision Aiding Process. R package (version 0.0.21). https://CRAN.R-project.org/package=MCDA

Ollerton, J., Winfree, R., Tarrant, S., 2011. How many flowering plants are pollinated by animals?. Oikos. 120, 321–326. https://doi.org/10.1111/j.1600-0706.2010.18644.x

Potts, S.G., Biesmeijer, J.C., Kremen, C., Neumann, P., Schweiger, O., Kunin, W.E. 2010. Global pollinator declines: trends, impacts and drivers. Trends Ecol. Evol. 25, 345–353. https://doi.org/10.1016/j.tree.2010.01.007

Raupp, M. J., Shrewsbury, P. M., Herms, D. A. 2010. Ecology of herbivorous arthropods in urban landscapes. Annual Rev. Entom. 55, 19–38. https://doi.org/10.1146/annurev-ento-112408-085351

R Core Team, 2019. R: A language and environment for statistical computing. R Foundation for Statistical Computing. https://www.R-project.org/ x(accessed 28 December 2021).

Robinson, N., 1993. Place and plant design–plant signatures. The Landscape. 53, 26–28.

Rodger, J.G., Bennett, J.M., Razanajatovo, M., Knight, T.M., van Kleunen, M., Ashman, T.L., Steet, J.A., Hui, C., Arceo-Gómez, G., Burd, M., Burkle, L.A., Burns, J.H., Durka, W., Freitas, L., Kemp, J.E., Li, J., Pauw, A., Vamosi, J.C., Wolowskijing, M., Xia, J., Ellis, A.G., 2021. Widespread vulnerability of flowering plant seed production to pollinator declines. Sci. Adv. 7, eabd3524. https://doi.org/10.1126/sciadv.abd3524

Rosasco, P., Perini, K., 2019. Selection of (green) roof systems: A sustainability-based multi-criteria analysis. Buildings. 9, 134. https://doi.org/10.3390/buildings9050134

Saaty, T.L., 1980. The Analytic Hierarchy Process. McGraw-Hill, New York.

Saaty, T.L., 1990. How to make a decision: the analytic hierarchy process. Eur. J. Oper. Res. 48, 9–26. https://doi.org/10.1016/0377-2217(90)90057-I

Saaty, T.L., 2004. “Decision making — the Analytic Hierarchy and Network Processes (AHP/ANP).” J. Syst. Sci. Syst. Eng. 13, 1–35. https://doi.org/10.1007/s11518-006-0151-5

Saaty, T.L., 2008. Decision making with the analytic hierarchy process. Int. J. Serv. Sci. 1, 83–98.

Saaty, T.L., De Paola, P., 2017. Rethinking design and urban planning for the cities of the future. Buildings. 7, 76. https://doi.org/10.3390/buildings7030076

Salman, I.N., Blaustein, L., 2018. Vegetation cover drives arthropod communities in mediterranean/subtropical green roof habitats. Sustainability. 10, 4209. https://doi.org/10.3390/su10114209

Schneider, C.A., Rasband, W.S., Eliceiri, K.W., 2012. NIH Image to ImageJ: 25 years of image analysis. Nat. Methods. 9, 671–675. https://doi.org/10.1038/nmeth.2089

Si, J., Marjanovic-Halburd, L., Nasiri, F., Bell, S., 2016. Assessment of building-integrated green technologies: A review and case study on applications of Multi-Criteria Decision Making (MCDM) method. Sustain. Cities Soc. 27, 106–115. https://doi.org/10.1016/j.scs.2016.06.013

Sikorski, P., Gawryszewska, B., Sikorska, D., Chormański, J., Schwerk, A., Jojczyk, A., Ciężkowski, W., Archiciński, P., Łepkowski, M., Dymitryszyn, I., Przybysz, A., Wińska-Krysiak, M., Zajdel, B., Matusiak, J., Łaszkiewicz, E., 2021. The value of doing nothing– How informal green spaces can provide comparable ecosystem services to cultivated urban parks. Ecosyst. Serv. 50, 101339. https://doi.org/10.1016/j.ecoser.2021.101339

Suárez, M.A., 2020. Avances en la introducción de cubiertas naturadas sustentables para zonas urbanas y periurbanas de la ciudad de Córdoba: selección de materiales de Glandularia spp. Doctoral dissertation, Universidad Católica de Córdoba, Córdoba. http://pa.bibdigital.ucc.edu.ar/2413/

Sutton, R.K., 2015. Introduction to green roof ecosystems, in: Sutton, R. (Ed.), Green Roof Ecosystems. Ecological Studies (Analysis and Synthesis), vol 223. Springer, Cham., pp. 1–25. https://doi.org/10.1007/978-3-319-14983-7_1

Thuring, C., Grant, G., 2016. The biodiversity of temperate extensive green roofs–a review of research and practice. Isr. J. Ecol. Evol. 62, 44–57. https://doi.org/10.1080/15659801.2015.1091190

Vlachokostas, C., Michailidou, A.V., Matziris, E., Achillas, C., Moussiopoulos, N., 2014. A multiple criteria decision-making approach to put forward tree species in urban environments. Urban Clim. 10, 105–118. https://doi.org/10.1016/j.uclim.2014.10.003

Wang, J.W., Poh, C.H., Tan, C.Y.T., Lee, V.N., Jain, A., Webb, E.L., 2017. Building biodiversity: drivers of bird and butterfly diversity on tropical urban roof gardens. Ecosphere. 8, e01905. https://doi.org/10.1002/ecs2.1905

Yakub, M., Tiffin, P., 2017. Living in the city: urban environments shape the evolution of a native annual plant. Glob. Change Biol. 23, 2082–2089. https://doi.org/10.1111/gcb.13528

Yee, E.G., Callahan, H.S., Griffin, K.L., Palmer, M.I., Lee, S., 2021. Seasonal patterns of native plant cover and leaf trait variation on New York City green roofs. Urban Ecosyst. 1–12. https://doi.org/10.1007/s11252-021-01134-2

Zhang, H., Lu, S., Fan, X., Wu, J., Jiang, Y., Ren, L., Wu, J., Zhao, H. (2021). Is sustainable extensive green roof realizable without irrigation in a temperate monsoonal climate? A case study in Beijing. Sci. Total Environ., 753, 142067. https://doi.org/10.1016/j.scitotenv.2020.142067

